# Context-dependent effects of *CDKN2A* and other 9p21 gene losses during the evolution of oesophageal cancer

**DOI:** 10.1101/2024.01.24.576991

**Authors:** Piyali Ganguli, Celia C. Basanta, Amelia Acha-Sagredo, Hrvoje Misetic, Maria Armero, Akram Mendez, Aeman Zahra, Ginny Devonshire, Gavin Kelly, Adam Freeman, Mary Green, Emma Nye, Anita Bichisecchi, Paola Bonfanti, Oesophageal Cancer Clinical, Molecular Stratification (OCCAMS) Consortium, Manuel Rodriguez-Justo, Jo Spencer, Rebecca C. Fitzgerald, Francesca D. Ciccarelli

## Abstract

*CDKN2A* is a tumour suppressor located in chromosome 9p21 and frequently lost in Barrett’s oesophagus (BO) and oesophageal adenocarcinoma (OAC). How *CDKN2A* and other 9p21 gene co-deletions affect OAC evolution remains understudied. We explored the effects of 9p21 loss in OACs and cancer progressor and non-progressor BOs with matched genomic, transcriptomic, and clinical data. Despite its cancer driver role, *CDKN2A* loss in BO prevents OAC initiation by counter-selecting subsequent *TP53* alterations. 9p21 gene co-deletions predict poor patient survival in OAC but not BO through context-dependent effects on cell cycle, oxidative phosphorylation, and interferon response. Immune quantifications using bulk transcriptome, RNAscope and high-dimensional tissue imaging showed that *IFNE* loss reduces immune infiltration in BO but not OAC. Mechanistically, *CDKN2A* loss suppresses the maintenance of squamous epithelium, contributing to a more aggressive phenotype. Our study demonstrates context-dependent roles of cancer genes during disease evolution, with consequences for cancer detection and patient management.

## BACKGROUND

*CDKN2A* is among the most frequently damaged cancer genes, with loss of function (LoF) reported in at least 35 different tumour types across 12 organ systems^1^. *CDKN2A* acts as a tumour suppressor by inducing cell cycle arrest and cellular senescence^2^ as well as preventing angiogenesis^3^, oxidative stress^4^, and metastasis^2^. Additionally, *CDKN2A* LoF predicts poor patient survival^5–7^.

*CDKN2A* LoF may occur through damaging point mutations, small indels or large deletions of chromosome 9p21.3 locus (hereon 9p21), an event observed in around 15% of cancers^8^. Depending on their length, 9p21 deletions may involve up to 26 genes, including other cell cycle regulators (*CDKN2B* and *KLHL9*), a metabolic enzyme (*MTAP*), and a cluster of 16 type-I interferons (**Figure 1A**). Recently, the loss of the whole locus, rather than *CDKN2A* alone, has been associated with poor survival and resistance to immunotherapy, possibly through the onset of an immune-cold tumour microenvironment (TME)^8^.

**Figure 1.**
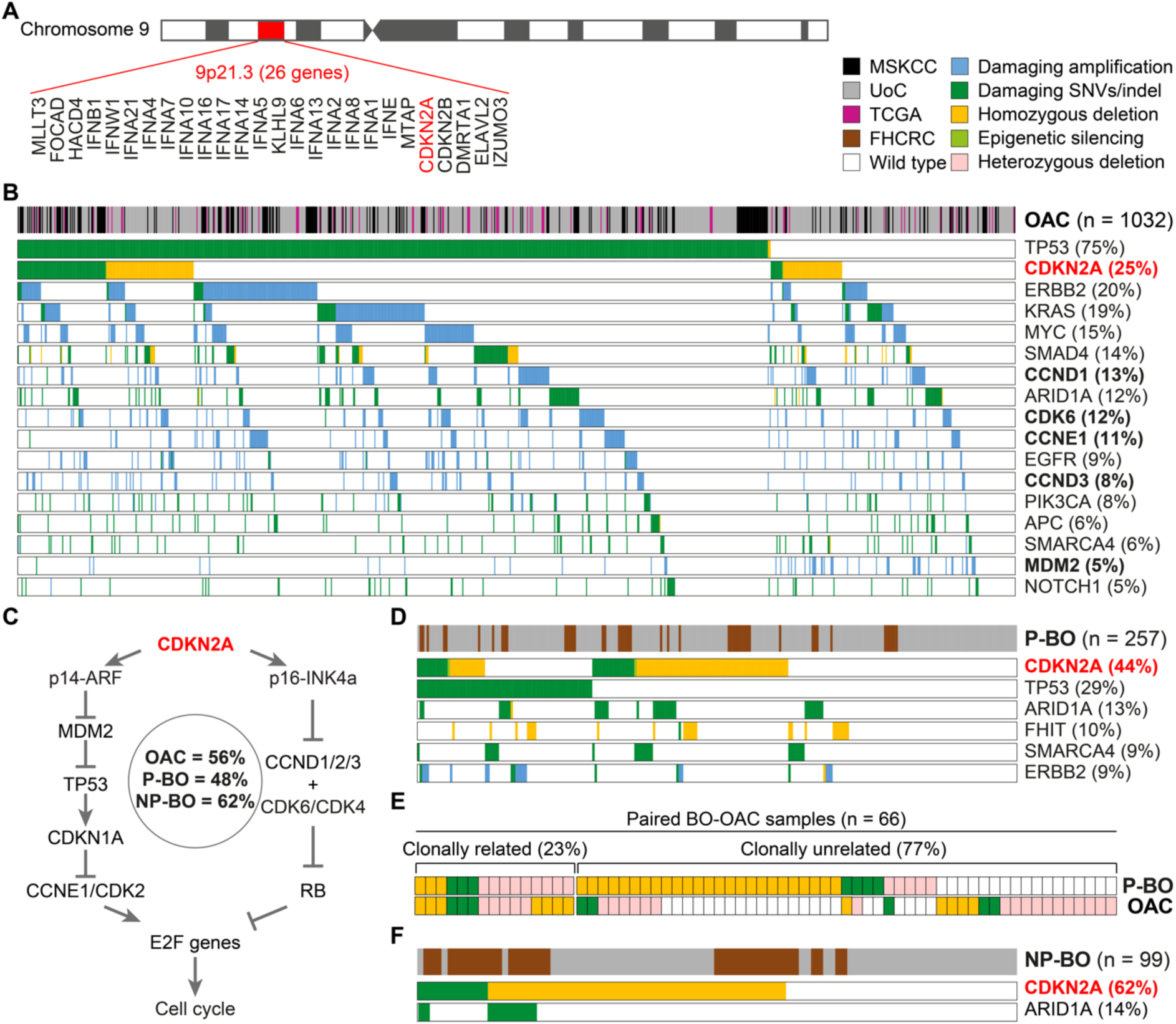
*CDKN2A* LoF occurrence in BO and OAC. **A.** Gene composition of chromosome 9p21 locus. **B.** Canonical OAC drivers damaged in at least 5% of 1032 OACs. All cell cycle regulators are reported in bold. **C.** Alterations in cell cycle regulators in BO and OAC. *CDKN2A* gene products (p14-ARF and p16-INK4a) regulate the cell cycle through the E2F genes^46^. p14-ARF blocks MDM2 and TP53 degradation, which induces *CDKN1A* transcription. CDKN1A in turn inhibits the CCNE1/CDK2 complex ultimately blocking cell cycle through E2F1 inhibition. p16-INK4a directly inhibits the CCND/CDK6/CDK4 complex preventing RB1 phosphorylation. Unphosphorylated RB1 can bind E2F1 leading to cell cycle arrest. *CDKN2A* LoF favours cell cycle progression resulting in uncontrolled cell proliferation. Values within the circle represent the proportion of OACs, P-BOs and NP-BOs with at least one damaged cell cycle regulator (except TP53). **D.** Canonical OAC drivers damaged in at least 5% of 257 P-BOs. **E.** Paired BO-OACs with *CDKN2A* LoF. Clonally related alterations refer to either identical *CDKN2A* alterations in both lesions or *CDKN2A* alterations in BO that could further evolve in OAC. **F.** Canonical drivers damaged in at least 5% of 99 NP-BO. Alteration frequency of OAC canonical drivers in (**B**, **D** and **F**) is indicated in brackets. The alteration frequency of all OAC drivers in the three cohorts is available in **Table S2**. FHCRC, Fred Hutchinson Cancer Research Center; LoF, loss of function; MSKCC, Memorial Sloan Kettering Cancer Center; NP-BO, non-progressor Barrett’s oesophagus; OAC, oesophageal adenocarcinoma; P-BO, progressor Barrett’s oesophagus; SNV, single nucleotide variant; TCGA, The Cancer Genome Atlas; UoC, University of Cambridge.

Dissecting the consequences of individual 9p21 gene losses is not straightforward because of their co-occurrence. Recently, the induction of different 9p21 deletions in pancreatic cancer mouse models enabled observation of reduced CD8^+^ T cell infiltration only when the IFN cluster was co-deleted with *CDKN2A*, *CDKN2B* and *MTAP*^9^. *IFNE*, one of 9p21 type-I interferons (**Figure 1A**), is a tumour suppressor in ovarian cancer^10^ and IFNE treatment promotes CD8^+^ T cell activation while reducing T regulatory cells (Tregs) and myeloid-derived suppressor cells (MDSCs)^10^. Also *MTAP* can regulate CD8^+^ and CD4^+^ T cell infiltration in melanoma mouse models by controlling methylthioadenosine accumulation in their TME^11^. All these studies started to unveil that at least some of the effects previously ascribed to *CDKN2A* LoF are in fact due to the loss of other 9p21 genes.

*CDKN2A* LoF has long been known as an early event in the evolution of oesophageal adenocarcinoma (OAC), occurring already in its precursor Barrett’s Oesophagus (BO)^12–16^. Consequently, *CDKN2A* LoF has been proposed to drive OAC initiation by favouring BO clonal selective sweeps and subsequent alterations of additional drivers, most frequently *TP53*^17–20^. Recently, this model has been replaced by an alternative one where early *TP53* LoF would enable whole genome doubling with consequent acquisition of additional drivers^21,22^. The role of *CDKN2A* LoF in OAC initiation remains controversial. Some studies reported significantly higher frequency of *CDKN2A* LoF in BOs progressing to OAC compared to BOs that did not progress^23–27^, implying that *CDKN2A* inactivation favours cancer initiation. Other studies found either no difference between progressor and non-progressor BOs^22,28–31^ or a higher frequency of *CDKN2A* LoF in non-progressor BOs^15^. This uncertainty raises questions on the role of *CDKN2A* in BO and OAC evolution. Moreover, very little is known about the function of the remaining 9p21 genes.

Here we investigated how the loss of *CDKN2A* and other 9p21 genes affects OAC initiation and progression. We compared genomic, transcriptomic, and survival data from large and clinically annotated cohorts of OAC and BO patients who progressed or did not progress to cancer. We validated the results *in vitro* and studied the effect of 9p21 loss on BO and OAC TME by high-dimensional tissue profiling coupled with RNAscope. Finally, we rebuilt the causal gene regulatory networks linking *CDKN2A* gene loss to specific downstream functional effects. Our results suggested that the same genetic alterations of *CDKN2A* and other 9p21 genes have different effects in different contexts and stages of OAC evolution, with possible implications in patient management.

## RESULTS

### *CDKN2A* LoF drives BO and OAC evolution but does not trigger OAC initiation

We collected whole genome (WGS), whole exome (WES) and gene panel sequencing data for 1032 OACs from the literature^6,32–38^ or sequenced *de novo* by the Oesophageal Cancer Clinical and Molecular Stratification (OCCAMS) Consortium (**Table S1**). Our cohort reflected OAC high male prevalence, with almost 9:1 male-to-female incidence ratio^39^ (**Table S1**). To ensure consistency, we annotated damaging mutations and copy number alterations in all datasets using the same approach (**Methods, Figure S1**). Because *CDKN2A* can be silenced also via epigenetic modifications, we analysed methylation data for a subset of OACs^32,40^ (**Table S1**). We then identified the damaged drivers in each sample using a curated list of 54 known (canonical) OAC drivers (**Table S2**). In agreement with previous studies^29,32,41^, *CDKN2A* was the second most frequently damaged OAC driver, with LoF in 25% of samples (**Figure 1B**). More than 56% of OACs (90% considering also *TP53*) had damaging alterations in other cell cycle regulators (**Figure 1C, Table S2**), suggesting that cell cycle disruption is key in OAC evolution but does not always involve *CDKN2A*.

Next, we measured the frequency of *CDKN2A* LoF in 257 BOs that progressed to high-grade dysplasia or OAC (P-BOs), again sequenced for this study or gathered from published datasets^15,40,42–44^ (**Table S1**). *CDKN2A* LoF occurred significantly more frequently in P-BO than OAC (p = 4×10^-9^, two-sided Fisher’s exact test, **Figure 1D**), suggesting that OAC does not always originate from a *CDKN2A*-damaged BO. To further investigate this, we analysed 66 matched OAC-BO pairs with *CDKN2A* LoF in BO or OAC (**Table S1**). Only 15 matched lesions had either identical or clonally related *CDKN2A* alterations (**Figure 1E**), confirming that *CDKN2A* LoF is not required for precancer to cancer transition. Interestingly, 28 OACs lost *CDKN2A* independently of the paired BOs (**Figure 1E**), suggesting that either OAC developed from a different *CDKN2A*-damaged BO clone or *CDKN2A* LoF was acquired after transformation.

Finally, we analysed 99 BOs that did not progress to high-grade dysplasia or OAC (NP-BOs)^15,40,43,44^ (**Table S1**). The frequency of *CDKN2A* LoF in NP-BO was even higher than P-BO and OAC (p = 3×10^-3^ and p = 3×10^-13^, respectively, two-sided Fisher’s exact test, **Figure 1F**). Moreover, while in OAC the dysregulation of cell cycle could occur through alterations of other genes, *CDKN2A* was the only cell cycle gene damaged in BO (**Figure 1D**). Therefore, unlike OAC, only *CDKN2A* LoF is relevant for BO evolution.

As observed previously^22,45^, P-BOs had significantly more damaged drivers than NP-BOs (p = 7×10^-6^, two-sided Fisher’s exact test, **Table S2**), indicating that OAC initiation requires several driver events, most frequently *TP53* complete loss. Given its high recurrence, we used *TP53* LoF to assess the role of *CDKN2A* LoF in OAC initiation calculating the odds of cancer progression based on the mutational status of *CDKN2A* and *TP53* in BO. As expected, the odds of cancer progression in BOs with *TP53* LoF was 1 irrespective of *CDKN2A* status (**Table S3**), confirming that *TP53* is a strong driver of OAC initiation. However, the odds of cancer progression in BOs with *CDKN2A* LoF and wild type *TP53* was lower than those of BOs with both wild type genes (0.58 and 0.72, respectively, **Table S3**). This suggested that an early occurrence of *CDKN2A* LoF in BO may reduce the likelihood of OAC initiation. To test this further, we compared two logistic regression models, one assuming a role in OAC initiation only for *TP53* LoF (model 1) and the other for both *TP53* and *CDKN2A* LoFs (model 2, **Methods**). Model 2 was a significantly better predictor of OAC initiation than model 1 (p = 0.01, ANOVA test), with expected occurrences of P-BOs with any status of *TP53* and *CDKN2A* perfectly matching the observed occurrences (**Table S3**). The negative β coefficient of *CDKN2A* in model 2 further confirmed that *CDKN2A* LoF may reduce risk of cancer progression (**Methods, Table S3**).

### Additional *TP53* loss reduces proliferation of *CDKN2A* LoF BO cells

Next, we set out to investigate how *CDKN2A* LoF in BO could prevent OAC initiation. Since the proportion of BOs with both *CDKN2A* and *TP53* LoF was significantly lower than that of BOs with *CDKN2A* LoF only (p = 0.05, two-sided Fisher’s exact test, **Figure 2A**), we hypothesised that negative selection might act on BO cells losing both genes. To test this hypothesis, we compared *CDKN2A* and *TP53* LoF clonality in 580 OACs with WGS or WES data, since clonality informs on when alterations are acquired during cancer evolution. Despite the well-known OAC intra-tumour heterogeneity^14^, *CDKN2A* or *TP53* LoFs were clonal in almost 70% of OACs (397/580), confirming that both alterations are early events. However, OACs with fully clonal *CDKN2A* LoF were significantly fewer than those with fully clonal *TP53* LoF (p = 0.001, two-sided Fisher’s exact test, **Figure 2B**), suggesting that overall*TP53* LoF tends to predate *CDKN2A* LoF. In support of this, *CDKN2A* LoF occurred before *TP53* LoF in only 6% of the 47 OACs with LoF alterations in both genes as compared to 38% where *TP53* LoF occurred before that of *CDKN2A* (**Figure 2C**), This confirmed that the subsequent loss of *TP53* in the presence of *CDKN2A* LoF is a rare event suggesting that it might be selected against.

**Figure 2.**
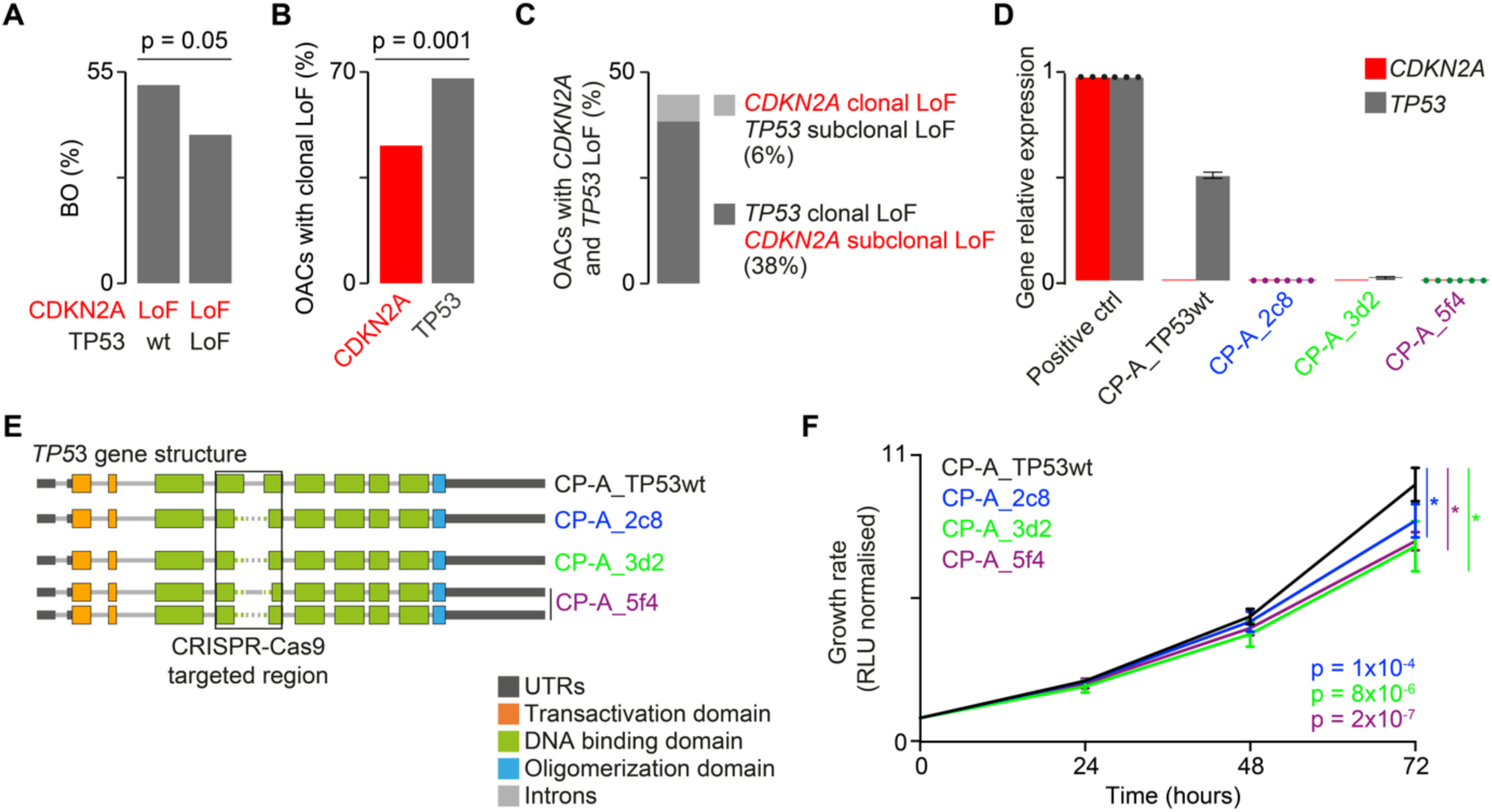
Effect of *TP53* loss in BO with *CDKN2A* LoF. **A.** Frequency of *CDKN2A* LoF in 356 BOs (257 P-BOs and 99 NP-BOs) with or without *TP53* LoF. Statistical significance was assessed using a two-sided Fisher’s exact test. **B.** Frequency of OACs with clonal LoF alterations in *CDKN2A* and *TP53* genes. For this analysis, 580/779 OACs with WGS or WES data and Lof in these genes were considered. Statistical significance was assessed using a two-sided Fisher’s exact test. **C.** Frequency of OACs with clonal and subclonal LoF alterations in *CDKN2A* and *TP53* genes in 47 OACs with WGS or WES data and damaging alterations in both these genes. **D.** *CDKN2A* and *TP53* gene expression levels quantified by RT–qPCR of RNA from *TP53* wild type CP-A cells (CP-A_ *TP53*wt), three *TP53* KO clones (CP-A_2c8, CP-A_3d2, CP-A_5f4) and control RNA relativised to *ACTB* expression. Mean and upper–lower limits of RQ are shown. **E.** *TP53* gene structure in CP-A_*TP53*wt, CP-A_2c8, CP-A_3d2, and CP-A_5f4 cells. Exon-intron arrangement was derived from the UCSF genome browser (https://genome.ucsc.edu/) based on NM_000546 mRNA sequence (chr17:7,668,421-7,687,490, hg38 assembly). Dotted lines represent edited regions. **F.** Growth curves of CP-A_*TP53*wt, CP-A_2c8, CP-A_3d2, and CP-A_5f4 cells. Proliferation was assessed every 24 h and normalised to time zero. Mean values at 72 h were compared by two-tailed Student’s t-test. Error bars show standard deviation. Three biological replicates were performed, each in two to four technical replicates. BO, Barrett’s oesophagus; ctrl, control; KO, knockout; LoF, loss of function; NP-BO, non-progressor Barrett’s oesophagus; OAC, oesophageal adenocarcinoma; P-BO, progressor Barrett’s oesophagus; RLU, relative light unit; RT–qPCR, real time quantitative PCR; RQ, relative quantification; UTR, untranslated region; wt, wild type.

Interestingly, BAR-T cells, derived from BO with constitutive loss of *CDKN2A*, increase cell doubling times upon *TP53* knock-down^47^, supporting the hypothesis that the additional loss of *TP53* reduces cell growth rate. To test this experimentally, we induced *TP53* knockout (KO) in metaplastic BO CP-A cells derived from a male individual with *CDKN2A* LoF and wild type *TP53*^48^. First, we confirmed that CP-A cells expressed *TP53* but did not express *CDKN2A* (**Figure 2D**). We then used CRISPR-Cas9 to edit *TP53* (**Table S4**) and performed single cell cloning to expand cell colonies. To control for off target effects and clonal differences, we selected three clones with a partial deletion of *TP53* exons 5 and 6 (**Figure 2E**), as assessed via amplicon sequencing (**Table S4**). We confirmed that these clones did not express *CDKN2A* nor *TP53* (**Figure 2D**). The fact that we could isolate clones losing both genes implied that BO cells with *CDKN2A* LoF can survive subsequent *TP53* loss. However, compared to *TP53* wild type CP-A cells, all three *TP53* KO CP-A clones showed significantly slower growth rate that was already visible after 72 hours (two-tailed t-test test, **Figure 2F**).

This was in line with the reported increase in cell doubling times of *TP53* knock-down BAR-T cells^47^ and supported the tumour preventive role of early *CDKN2A* inactivation due to the reduced fitness, defined as proliferative capacity, of cells additionally losing *TP53*.

### LoF of 9p21 genes predicts poor survival in OAC but not in BO

Since *CDKN2A* LoF has been associated with poor patient survival^5–7^, we investigated the survival effect of *CDKN2A* and other 9p21 gene LoF in our extended BO and OAC cohorts. OAC patients with *CDKN2A* LoF showed significantly worse survival than those with the wild type gene (**Figure 3A**). This held true even when patients with *CDKN2A* homozygous deletions (**Figure 3B**) or damaging mutations (**Figure 3C**) were considered separately. However, we did not observe lower survival in patients with *CDKN2A* heterozygous deletions only (**Figure S2A**), suggesting that *CDKN2A* complete loss is required to affect prognosis. Damaging alterations in *TP53* or other cell cycle regulators had no effect on survival (**Figure S2B**-**F**) despite their frequent OAC alterations (**Figure 1C**). Therefore, the survival effect of *CDKN2A* LoF does not depend on its function as cell cycle regulator. Moreover, *CDKN2A* LoF was not a predictor of worse survival in P-BO (**Figure 3D**), again suggesting context-dependent consequences of its loss.

**Figure 3.**
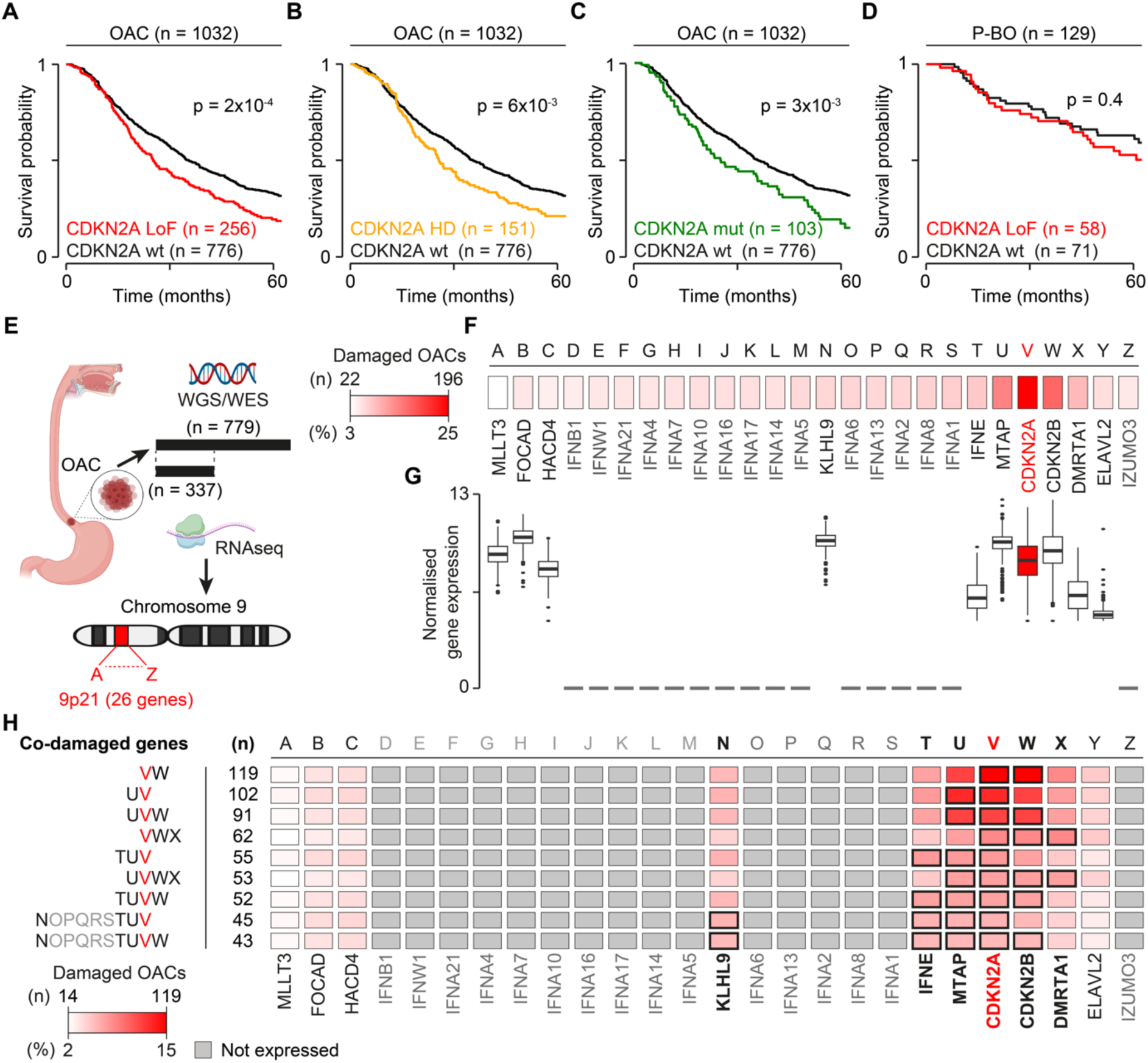
Effect of the LoF of *CDKN2A* and other 9p21 genes on survival. Kaplan-Meier survival curves of OAC patients with wild type *CDKN2A* compared to those with all types of LoFs (**A**), only homozygous deletions (**B**), and only LoF mutations (**C**). **D.** Kaplan-Meier survival curves of P-BO patients with and without *CDKN2A* LoF. **E.** Approach to test the effect of the co-damage in 9p21 genes on survival of OAC patients. Only 779 OACs with WGS or WES data were used for the survival analysis while 337 of them with RNAseq data were used to measure 9p21 gene expression. Letters correspond to the 26 genes according to their order in the chromosomal locus. **F.** LoF frequency of 9p21 genes in 779 OACs. **G.** Distribution of normalised expression values in the 9p21 genes in 337 OACs. **H.** Kaplan-Meier survival analysis of OAC patients with co-alterations in the ten expressed 9p21 genes and 413 OAC patients with a wild type locus. Only groups with significantly poor survival (FDR <0.1) are shown and genes of interest are circled in black. All groups used in the analysis are listed in **Table S5**. The minimum and maximum number and percent of damaged OACs in (**F**) and (**H**) are reported in the corresponding heatmap. HD, homozygous deletion; LoF, loss-of-function; OAC, oesophageal adenocarcinoma; P-BO, progressor Barrett’s oesophagus; WES, whole exon sequencing; WGS, whole genome sequencing; WT, wild type. Cartoon in **E** was created with BioRender.com.

We then investigated whether the co-occurring loss of other 9p21 genes could also contribute to poor survival, restricting the analysis to 779 OACs with WGS or WES data (**Figure 3E**). Although *CDKN2A* was the most frequently occurring alteration in the locus, confirming that it is the event under positive selection, the other 25 genes were frequently co-lost with it (**Figure 3F**). However, only ten 9p21 genes were expressed in OAC (**Figure 3G**) or normal oesophagus (**Figure S3**), suggesting that the loss of the remaining 16 genes likely had no functional consequences. We therefore tested the potential impact on survival of the ten 9p21 expressed genes by dividing OAC patients in nine groups. Each of these groups represented at least 5% of the cohort and was composed of patients with the same 9p21 mutation and copy number profile (**Table S5**). Patients in all nine groups had worse survival than 413 OAC patients with a wild type 9p21 locus (FDR <0.1, **Figure 3H, Table S5**). All patients lost *KLHL9, IFNE*, *MTAP*, *CDKN2A*, *CDKN2B* and *DMRTA1* (**Figure 3H**), suggesting that alterations in these genes may contribute to poor prognosis.

### LoF of 9p21 genes has distinct functional consequences in BO and OAC

Our results suggested that the LoFs of *CDKN2A* and other 9p21 genes have functional and survival consequences that depend on time and context. Disentangling these variable effects is challenging because 9p21 genes are often co-damaged (**Figure 3F**). To tease out the contribution of individual 9p21 genes, we divided 22 NP-BOs, 108 P-BOs and 337 OACs with matched genomic and transcriptomic data (**Table S1**) into four groups (**Figure 4A**). Each group had the same LoF profile of the six genes whose loss impacted survival (*KLHL9, IFNE*, *MTAP*, *CDKN2A*, *CDKN2B* and *DMRTA1*, **Figure 3H**). Group 1 included all samples with *CDKN2A* LoF independently of the status of the other genes (**Figure 4B**), closely resembling the cohorts tested in the survival analysis (**Figure 3A and 3D**). The other three groups were subsets of group 1 with variable LoF frequency of in the six genes (**Figure 4B**).

**Figure 4.**
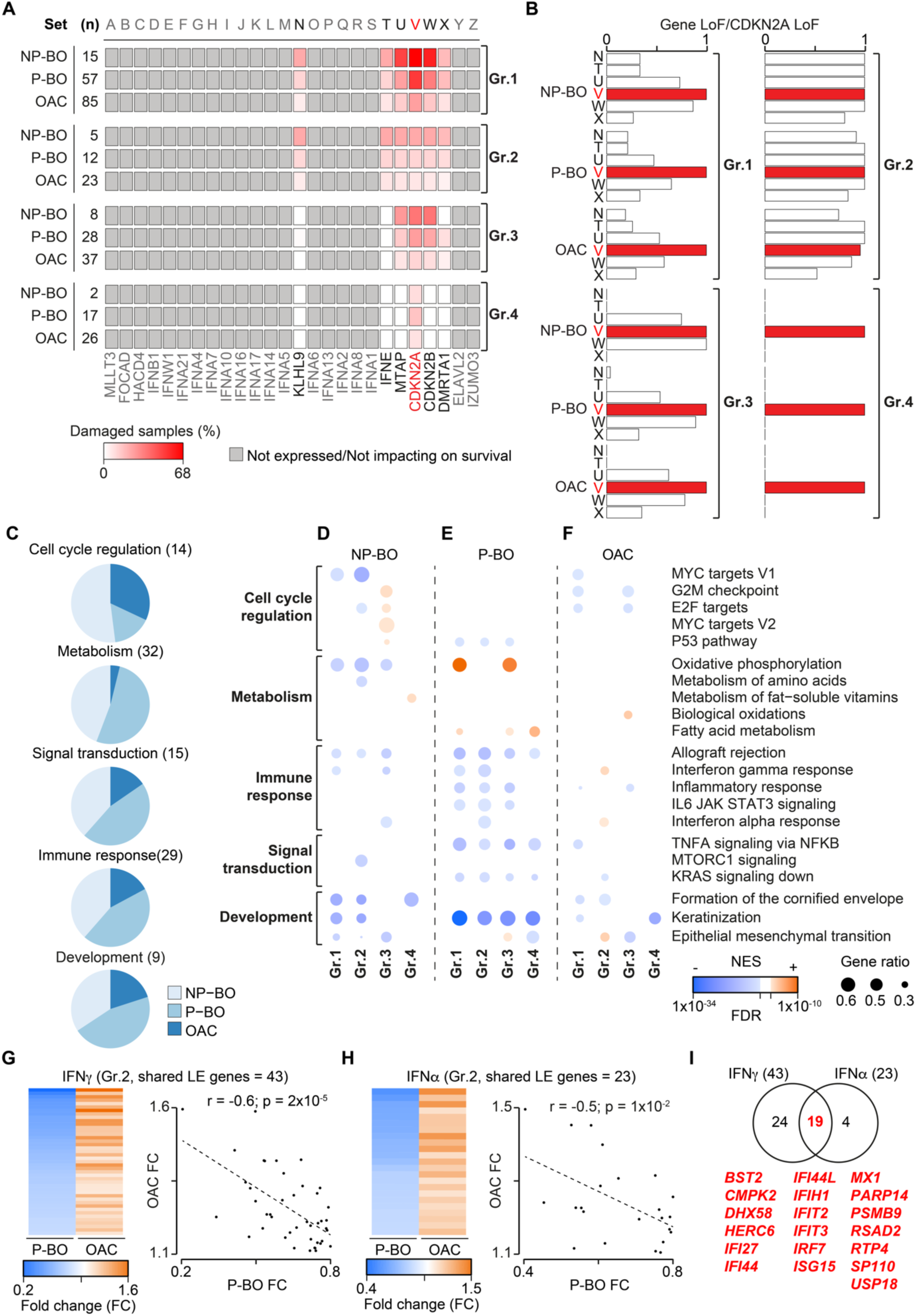
Functional consequences of 9p21 gene LoF in BO and OAC. **A.** Frequency of damaged and damaged 9p21 genes in the four groups of NP-BOs, P-BOs estimated over 22 NP-BOs, 108 P-BOs and 337 OACs with matched genomic and transcriptomic data. **B.** Proportions of samples with LoF in *KLHL9* (N), *IFNE* (T), *MTAP* (U), and *CDKN2B* (W) and *DMRTA1* (X) over samples with *CDKN2A LoF* (V) in each group of NP-BOs, P-BOs and OACs. The number of samples in each group and condition is reported. **C.** Relative proportion of dysregulated pathways in NP-BO, P-BO and OAC cohorts mapping to cell cycle regulation, metabolism, signal transduction, immune response, and development. Number in brackets represent the number of unique pathways. Results of pre-ranked GSEA^50^ showing the NES, FDR and gene ratio (number of leading-edge genes over the total expressed genes) of pathways dysregulated in each group of NP-BOs **(D),** P-BOs **(E)** and OACs **(F)**. NES >0 indicates pathway upregulation while NES <0 indicates downregulation. Fold change of expression and correlation plot of the shared leading-edge genes of interferon gamma **(G)** and alpha **(H)** response pathways enriched in P-BO and OAC group 2 as compared to 9p21 wild type samples. Pearson’s correlation coefficients and associated p-values are reported for both pathways. **I.** Overlap of leading-edge genes between interferon gamma and alpha response pathways enriched in P-BO and OAC group 2. The 19 shared genes are listed. FC, Fold change; FDR, False Discovery Rate; LoF, loss of function; NES, Normalised enrichment score; NP-BO, non-progressor Barrett’s oesophagus; OAC, oesophageal adenocarcinoma; P-BO, progressor Barrett’s oesophagus.

We identified the dysregulated biological processes in each group as compared to the corresponding 9p21 wild type samples by performing a pre-ranked gene set enrichment analysis (GSEA)^50^ in NP-BOs, P-BOs, and OAC separately. Overall, we detected 72, 62 and 28 unique pathways significantly dysregulated (FDR ≤0.01) in NP-BO, P-BO and OAC, respectively (**Table S6**). Almost 80% of these pathways mapped to only five biological processes, namely cell cycle regulation, metabolism, immune response, signal transduction, and development. Overall NP-BO and P-BO showed a higher fraction of dysregulated pathways than OAC (**Figure 4C**), suggesting that 9p21 LoF had higher impact in pre-malignant conditions.

As expected, given *CDKN2A*, *CDKN2B* and *KLHL9* role in cell cycle regulation role, we found cell cycle dysregulation across groups and conditions except group 4 (*CDKN2A* LoF only; **Figure 4D-F, Table S6**), suggesting that the co-deletion of *KLHL9, CDKN2A* and *CDKN2B* maximises the effect.

*CDKN2A* LoF alone might not be sufficient also to trigger metabolic or immune dysregulation (**Figure 4D-F, Table S6**). In this case *MTAP* and *IFNE* LoF could play a role given their functions in metabolic reprogramming^51,52^ and activation of immune response through metabolic regulation^53^, respectively. Interestingly, oxidative phosphorylation was consistently downregulated in NP-BO, upregulated in P-BO, and showed no difference in OAC (**Figure 4D-F, Table S6**). This once again suggested that the same genetic alterations may trigger different functional responses depending on the context. Similarly, the disruption of immune pathways differed between BO and OAC (**Figure 4D-F, Table S6**). While interferon alpha and gamma responses were consistently downregulated in NP-BO and P-BO, both were upregulated in OAC, particularly in group 2 (**Figure 4B**). Consistently, we observed a significant inverse correlation between expression fold changes of interferon gamma (**Figure 4G**) and alpha (**Figure 4H**) genes in BO and OAC groups 2 compared to 9p21 wild type samples. Moreover, there was substantial overlap between altered genes in the two pathways (**Figure 4I**), suggesting a comprehensive transcriptional reprogramming of interferon response. The most likely candidates for this reprogramming were again *MTAP*, given its recently reported ability to regulate the TME^11^, and *IFNE*, a type-1 interferon expressed in adult epithelia. Since the effect was most visible in group 2, which had LoF in both genes, and not in group 3, which had *MTAP* LoF and *IFNE* wild type (**Figure 4A,B**), the effect on interferon response might be due to *IFNE* loss.

*CDKN2A* LoF alone might instead be enough for the pervasive downregulation of keratinisation genes given that these pathways were consistently dysregulated also in group 4 (**Table S6, Figure 4D-F**).

### Loss of *IFNE* reduces immune infiltration in BO but not OAC

To further investigate the opposite effect of *IFNE* on interferon alpha and gamma response in BO and OAC (**Figure 4G,H**), we quantified the infiltration of 18 immune cell populations in NP-BOs, P-BOs and OACs from their bulk transcriptomic data. We then compared the abundance of immune infiltrates between each of the four 9p21 LoF groups (**Figure 4A**) and the corresponding 9p21 wild type samples.

Immune infiltrates were depleted in NP-BO groups 1-3 (**Figure 5A, Table S7**) and P-BO groups 1 and 2 as compared to 9p21 wild type samples (**Figure 5B, Table S7**), where the impact of *IFNE* LoF was more appreciable. This again suggested that the immune depletion is a consequence of *IFNE* loss consistent with recent observations of a cold TME when *IFNE*^10^ or the whole IFN locus^9^ are lost in melanoma ovarian, or pancreatic cancers (**Table S8**). However, the same studies also reported an increased infiltration of T regs, MDSCs and B cells (**Table S8**) that we did not observe (**Figure 5A, B**). The TME of group 4 (*CDKN2A* LoF only) was not significantly different to that of 9p21 wild type samples in both NP-BO and P-BO, confirming that *CDKN2A* LoF does not directly interfere with the immune system.

**Figure 5.**
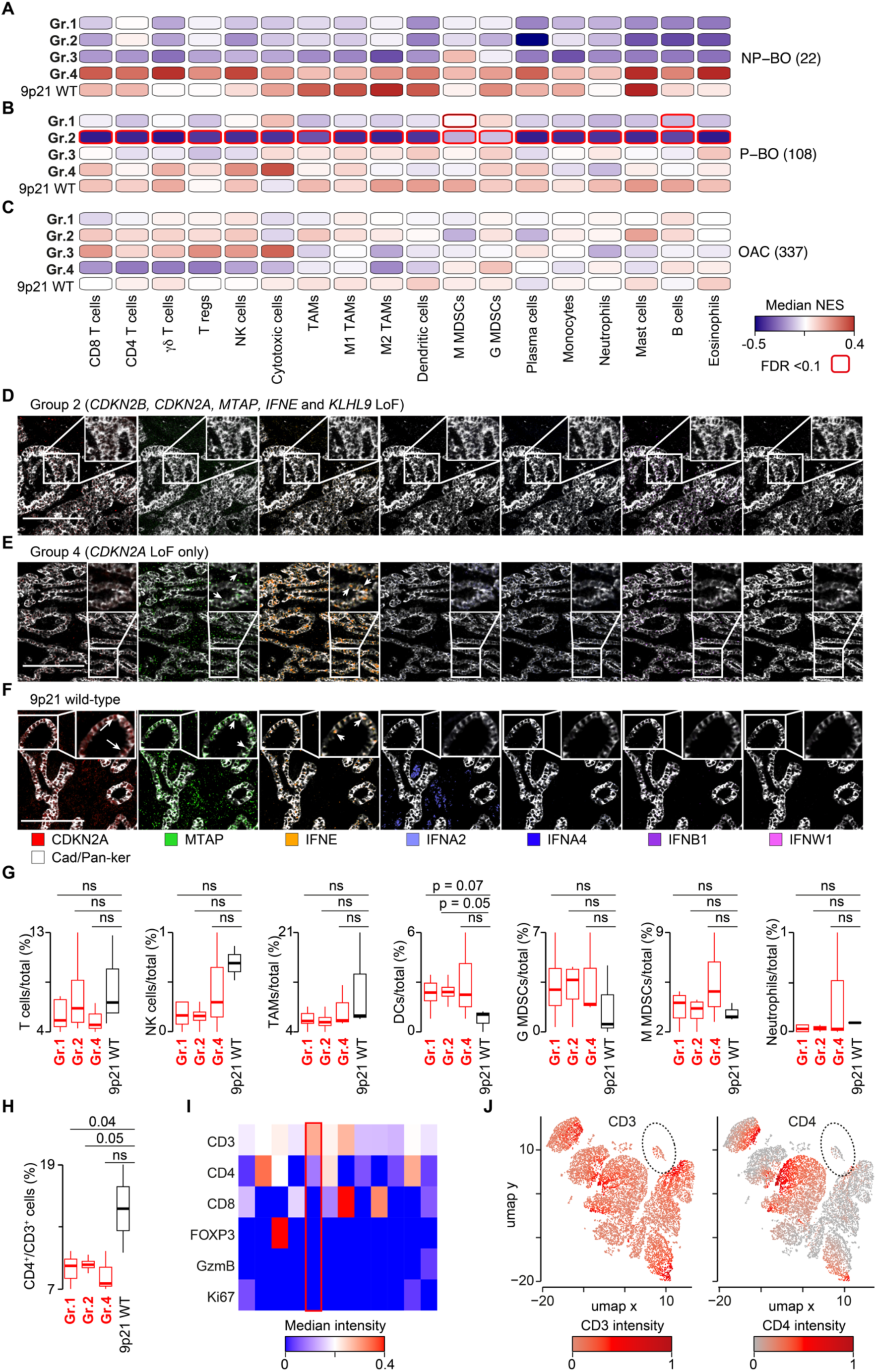
Impact of 9p21 gene loss on immune infiltration in BO and OAC. Comparison of NES scores of 18 immune populations between 9p21 LoF and wild type samples in NP-BOs **(A)**, P-BOs **(B)** and OACs **(C)**. NES distributions were compared using a two-sided Wilcoxon’s rank sum test and corrected for multiple testing using the Benjamini– Hochberg method. Number of samples are reported in brackets. Immune populations with significant differences (FDR < 0.1) are circled. Representative IMC images from group 2 **(D)**, group 4 **(E)** and 9p21 wild type **(F)** OACs showing the expression of 9p21 targeted proteins and mRNAs. Cadherin-1 and pan-keratin denote tumour. Arrows indicate examples of epithelial staining Scale bar: 200 μm. **G**. Relative abundance of immune cells over all cells in 9p21 LoF and wild type OACs. Samples in groups 2 and 4 were pooled together to form group 1 (n = 7). Distributions were compared using a two-sided Wilcoxon rank sum test. **H.** Relative abundance of CD4^+^ cells over all CD3^+^ cells in 9p21 LoF and wild-type OACs. Distributions were compared using a two-sided Wilcoxon rank sum test. **I.** Median marker intensity across the T cell clusters at a clustering resolution of 0.5. **J.** UMAP map of 9750 T cells in ten OACs. Cells were grouped in 12 clusters based on the expression of six markers and coloured according to the mean intensities of CD3 and CD4. The cluster enriched in group 1 is circled. FDR, False Discovery Rate; LoF, loss of function; NES, normalised enrichment score; NP-BO, non-progressor Barrett’s oesophagus; P-BO, progressor Barrett’s oesophagus; OAC, oesophageal adenocarcinoma.

Unlike other cancer types (**Table S8**) and BO (**Figure 5A, B**), we did not observe any significant TME difference between 9p21 LoF and wild type OACs (**Figure 5C, Table S7**). To investigate this at higher resolution, we performed high-dimensional imaging mass cytometry (IMC) on tissue sections representative of group 1, group 2, group 4 and 9p21 wild type OACs (**Table S9**). We used a panel of 26 antibodies targeting structural, immune, and 9p21-encoded proteins as well as RNAscope probes against *IFNE* and *IFNB1* mRNAs to increase the detection signal (**Table S10**). We confirmed that group 2 lost the expression of all 9p21 encoded proteins in the tumour, while group 4 lost CDKN2A only compared to 9p21 wild type OACs (**Figure 5D-F**). Moreover, IFNE was the only interferon clearly expressed in OAC epithelium (**Figure 5D-F**).

We performed single cell segmentation of the IMC images to quantify T cells, NK cells, macrophages, dendritic cells, monocytic (M) and granulocytic (G) MDSCs, and neutrophils (**Methods**). We then compared the relative abundance of each immune population over all cells in each slide across OAC groups. We confirmed no significant difference in immune infiltration between 9p21 LoF and wild type OACs, except for a borderline significant enrichment in dendritic cells in groups 1 and 2 (**Figure 5G**). We further applied unsupervised clustering to T cells and macrophages, for which we had multiple markers (**Table S10**), to test whether there was any difference in specific subpopulations. Again, we detected no major differences in any subpopulations of macrophages or T cells, except a borderline significant depletion of CD4^+^ T cells in groups 1 and 2 compared to 9p21 wild type OAC (**Figure 5H-J**). These results confirmed that, unlike BO, the loss of *IFNE* or any other 9p21 genes does not lead to any major difference in the TME of OAC.

### *CDKN2A* LoF favours squamous to columnar epithelium transition

We observed a pervasive downregulation of processes responsible for terminal differentiation of keratinocytes, such as keratinisation and formation of the cornified envelope, across all 9p21 LoF groups **(Figure 4D-F**). In particular, P-BO and OAC groups 4 were associated with the downregulation of keratinisation, suggesting that *CD2KNA* LoF alone was sufficient for triggering this process. To gain further mechanistic insights, we rebuilt the gene regulatory network linking *CD2KNA* LoF to keratinisation in P-BO and OAC group 4 (**Figure 6A**).

**Figure 6.**
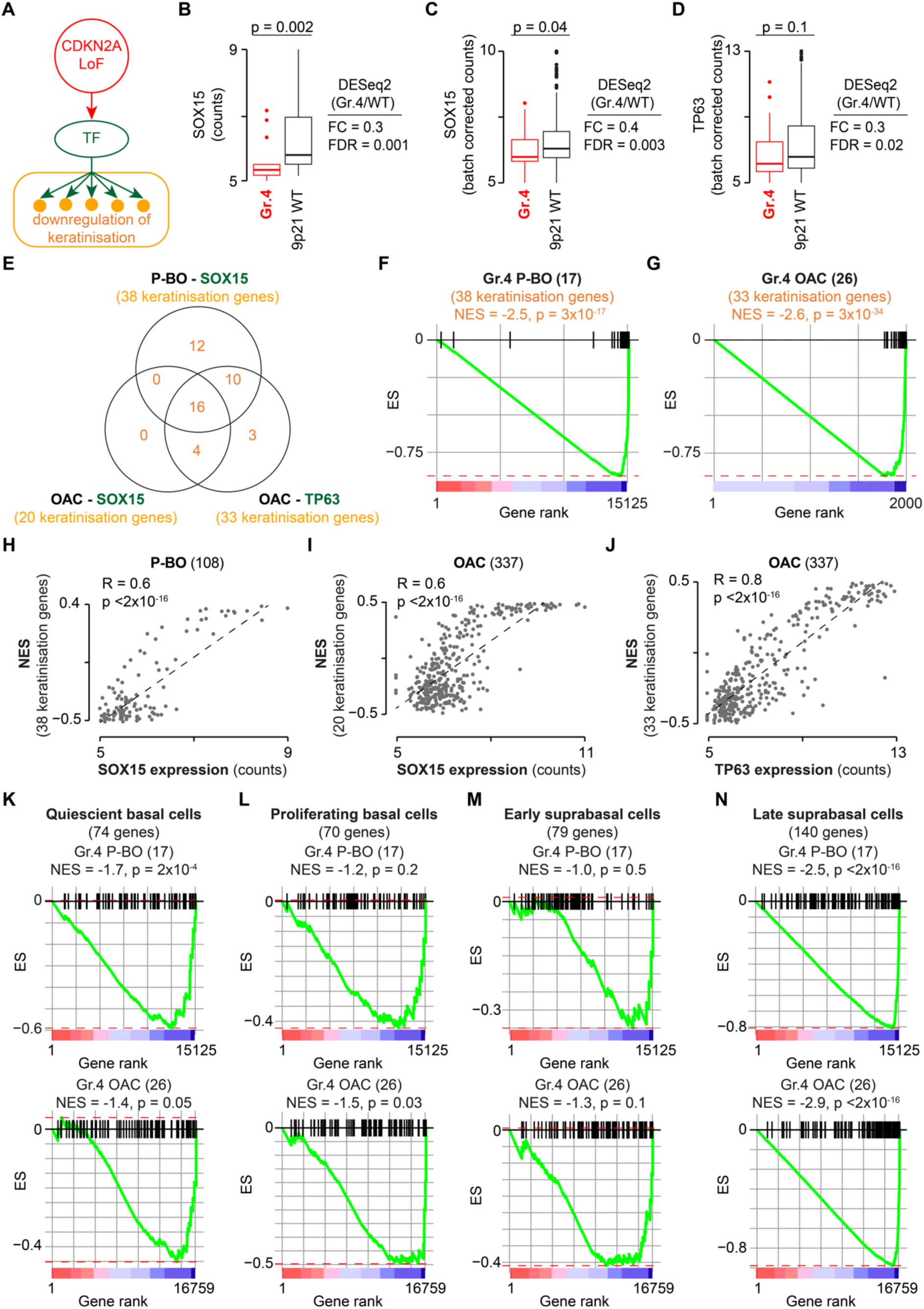
Impact of *CDKN2A* LoF on epithelium differentiation in P-BO and OAC. **A.** Gene regulatory network linking *CDKN2A* LoF to the downregulation of keratinisation genes through TF deregulations. Distributions of gene expression values of *SOX15* in P-BO **(B)** and *SOX15* **(C)** and *TP63* **(D)** in OAC in group 4 and 9p21 wild-type samples. Distributions were compared using two-sided Wilcoxon’s rank sum test. FC and FDR from the differential gene expression analysis with DESeq2^58^ are also shown. **E.** Overlap between keratinisation genes targeted by *SOX15* and *TP63* in P-BO and OAC. Pre-ranked GSEA plots using as signatures keratinisation genes targeted by *SOX15* in P-BO **(F)** and by *SOX15* and *TP63* in OAC **(G)**. Genes were ranked from the most upregulated to the most downregulated in group 4 compared to 9p21 wild type samples. For OAC, only the top 2000 downregulated genes are shown. Correlation plots between keratinisation GSEA NES and the gene expression values of *SOX15* in P-BO **(H)** and *SOX15* **(I)** and *TP63* **(J)** in OAC. Spearman’s correlation coefficient and associated p-values are shown. Pre-ranked GSEA plots using gene signatures for quiescent basal cells **(K)**, proliferating basal cells **(L)**, early supra-basal cells **(M)** and late supra-basal cells **(N)** in P-BO and OAC group 4. LoF, loss of function; NES, normalised enrichment score; OAC, oesophageal adenocarcinoma; P-BO, progressor Barrett’s oesophagus; TF, transcription factor; GSEA, gene set enrichment analysis.

Using a three-step protocol (**Figure S4, Methods**), we identified eight and 14 causal models in P-BO and OAC, respectively, linking *CDKN2A* LoF directly to keratinisation gene downregulation through the perturbation of two TFs (SOX15 and TP63, **Table S11**). We further confirmed that these TFs were significantly downregulated in P-BO (**Figure 6B**) and OAC (**Figure 6C, D**) groups 4 as compared to 9p21 wild type samples. Overall, the gene modules controlled by SOX15 and TP63 included 45 keratinisation genes (**Table S11**), 16 (36%) of which were shared across all gene modules and 30 were shared between SOX15 and TP63 (**Figure 6E**). Therefore, the downregulation of these two TFs in *CDKN2A* LoF samples led to a comprehensive downregulation of the keratinisation transcriptional programme, as confirmed by a pre-ranked GSEA^50^ using keratinisation gene-derived signatures in P-BO (**Figure 6F**) and OAC (**Figure 6G**). Moreover, *SOX15* and *TP63* gene expressions were positively correlated with the enrichment score of the keratinisation genes **(Figures 6H-J)**, again confirming that the two TFs control their expression.

SOX15 regulates transcription of a large number of genes specific to oesophageal epithelium^54^ and TP63 is essential for development and maintenance of all stratified epithelia^55^. The transition from oesophageal squamous epithelium to intestinal columnar epithelium is a key feature in the initiation of BO and OAC^56^. Our data suggested that *CDKN2A* LoF leads to a downregulation of the transcriptional programme responsible for the maintenance of the squamous epithelium more robust and persistent than in *CDKN2A* wild type samples. Although this did not prove a direct causative role of *CDKN2A* LoF, it showed correlation between the two events. To further test the link between *CDKN2A* LoF and suppression of squamous epithelium, we performed pre-ranked GSEA^50^ using four independent gene signatures characteristic of cells composing the oesophageal epithelium, namely quiescent basal cells, proliferating basal cells, early supra-basal cells and late supra-basal cells^57^. We observed global downregulation of all four signatures in OAC and quiescent basal cells and late supra-basal cells in P-BO (**Figure 6K-N**). These results supported our hypothesis that *CDKN2A* LoF exacerbates a phenotype typical of OAC and this may contribute to more aggressive tumours.

## DISCUSSION

In this study, we dissected the role of *CDKN2A* and other 9p21 genes in OAC evolution, from the transformation of premalignant BO to the impact on patient survival.

Despite being an OAC driver, the early loss of *CDKN2A* has a tumour suppressive role supported by its higher occurrence in NP-BO than P-BO and OAC. This is consistent with other drivers whose alterations are more frequent in normal tissues than cancer, including *ERBB2*, *ERBB3*, *KRAS*, and *NOTCH1*^49^. The anti-tumorigenic function of *NOTCH1* is exerted through an increased fitness of *NOTCH1* mutant cells that outcompete early tumours^59^. For *CDKN2A* we propose a different mechanism whereby *TP53* mutations reduce the proliferative capacity of *CDKN2A* mutant BO cells that are therefore counter-selected. Since *TP53* loss is a strong driver of OAC initiation, the decrease of its occurrence induced by *CDKN2A* LoF also decreases tumour initiation. Recent studies observed tumour formation upon induction of *TP53* and *CDKN2A* double KO in mouse or human gastroesophageal organoids^60–62^. However, in these studies *TP53* and *CDKN2A* inactivation was induced concomitantly, *i.e.* targeting both genes at the same time. However, in real pre-cancer conditions, such as BO, mutations are acquired over time and cells with different genetic makeup and fitness coexist and compete for nutrient and space. Our results confirmed that the order of mutations is key to decide the fate of mutant cells in the initial phases of tumour evolution^49^.

It is tempting to speculate that the tumour preventive role of early *CDKN2A* LoF could be further developed as a novel marker of favourable prognosis in non-dysplastic BO. Endoscopic surveillance of BO is an integral component of the current OAC prevention paradigm, but the rate of progression to OAC is only 0.54/100 patient-years^63^. Identifying BOs with a lower risk of progression could substantially improve patient management, decreasing the burden of endoscopy for patients who have low chances to develop cancer.

*CDKN2A* LoF is the most frequent event in 9p21 locus, implying that the co-occurring loss of other 9p21 genes is due to genetic hitchhiking, with variable effects on cell cycle, oxidative phosphorylation, and interferon response depending on the stage and context of BO and OAC evolution. Most notably, *IFNE* exerts a tumour-suppressive role in BO but not OAC by reducing IFN response and inducing a cold immune microenvironment. Despite several reports of a lower infiltration of immune cells in cancers with reduced *CDKN2A* expression^64,65^, *CDKN2A* LoF alone does not change the immune composition of BO or OAC TME. This may be due to tumour-specific effects or to the fact that at least some cancer promoting roles previously attributed to *CDKN2A* LoF are in fact triggered by the loss of other 9p21 genes.

The association of *CDKN2A* LoF with bad prognosis is also context dependent and detectable only in OAC patients. It appears unrelated to the role of *CDKN2A* in cell cycle since alterations in other cell cycle regulators can drive OAC without affecting survival. A contribution towards a more aggressive OAC phenotype is likely due to a combination of effects, including the pervasive suppression of transcriptional programmes responsible for the maintenance of squamous epithelium. Although this is a common feature of BO and OAC^56^, it is significantly more pronounced when *CDKN2A* is lost and is achieved through *TP63* and *SOX15* downregulation. This could be an indirect effect of *CDKN2A* LoF on the E2F transcriptional programme since iASPP, which controls *TP63* expression^66^, is a target of E2F1^67^ and *SOX15*, in turn, is a target of TP63^68^.

Our study introduces the intriguing concept that the functional consequences of alterations in cancer genes may change during the evolution of disease, from preventing cancer transformation in the premalignant setting to favouring a more aggressive disease at later stages. This fits the emerging scenario whereby the functional consequences of cancer alterations and the fitness provided to the mutant cell are not invariable but depend on the cell genetic background^69^, neighbourhood^59^, or order of events as we showed here. If proven of general applicability, this may lead to a paradigm shift with consequences on the understanding and treatment of cancer.

## METHODS

### Sample collection and ethical approval

Single Nucleotide Variants (SNVs), indels and copy number (CN) data for 1032 primary OACs were collected from published studies and *de novo* sequenced samples (**Table S1**). In particular, WGS from 706 OACs was performed at the University of Cambridge (UoC, EGAD00001011191 and EGAD00001006083, https://ega-archive.org/). WES for 73 TCGA OACs were downloaded from the Genomic Data Commons portal (https://portal.gdc.cancer.gov/). Damaged genes for 253 Memorial Sloan Kettering Cancer Center (MSKCC) OACs that underwent targeted re-sequencing of 528^37^, 477^6^ and 970^38^ genes were downloaded from the cBioPortal (https://www.cbioportal.org/). In cases of multiple samples per patient, the sample with *CDKN2A* LoF was retained. Clinical data for the TCGA and MSKCC cohorts were obtained from the same sources. For the UoC cohort clinical data were derived from LabKey (https://occams.cs.ox.ac.uk/labkey). Bulk RNAseq data were available for 337 OACs, all of which had matched WGS or WES (**Table S1**). Of these, 264 were sequenced at the UoC (EGAD00001011190) and 73 derived from TCGA. Methylation data were available for 256 OACs (EGAD00010001822^40^ and TCGA^32^, **Table S1**).

WGS, WES and clinical data for 356 BOs were obtained from UoC (EGAD00001011191 and EGAD00001011189, which also includes samples from^43^ and^44^) and from the Fred Hutchinson Cancer Research Center (FHCRC)^15,42^ (**Table S1**). As for OAC, in cases of multiple samples per patient, the sample with *CDKN2A* LoF was retained. BOs were classified as progressors (P-BO, 257) and non-progressors (NP-BO, 99) based on whether patients progressed or not to high-grade dysplasia or OAC in a follow-up period of up to 17 years (**Table S1**).

Paired WGS BO and OAC data were available for 86 cases (EGAD00001011191 and EGAD00001006083, which also include samples from^34,35,43^). Methylation data for 57 BOs were derived from UoC (EGAD00010001838^40^ and EGAD00010001972 Katz-Summercorn, 2022 #72}). Bulk RNAseq data for 108 P-BOs and 22 NP-BOs were sequenced at the UoC (EGAD00001011190, including samples from^43^) (**Table S1**).

BO and OAC patients whose samples from UoC sequenced for this study were consented to research for the OCCAMS project (REC: 10/H0305/1 & IRAS:15757). Samples were collected either at endoscopy, staging laparoscopy, endoscopic mucosal resection, or surgical resection then snap frozen in liquid nitrogen. Samples were then embedded in optimal cutting temperature media for cutting of 1 x 3µM slide to be H&E stained and reviewed by a pathologist. Only tumour samples of >50% cellularity and BO samples with high intestinal metaplasia content proceeded to sequencing.

### DNA and RNA extraction, library preparation and variant calling

DNA and RNA were extracted using Qiagen AllPrep Mini kits, using a Precellys for tissue dissociation after all excess OCT was removed. Extracted nucleic acids were quantified by Qubit. Libraries were then prepared using Illumina PCR Free methods and sequenced on HiSeq 4000 or NovaSeq platforms. Paired-end whole genome sequencing at 50X target depth for OACs, P-BOs and NP-BOs and 30X target depth for matched normal (blood) was performed by Illumina, the Sanger Institute, or the CRUK Cambridge Institute on Illumina platforms. Quality checks were performed using FastQC (http://www.bioinformatics.babraham.ac.uk/projects/fastqc/). For mutation calling, sequencing reads were aligned against the reference genome (hg19/GRCh37) using BWA-MEM^70^. Aligned reads were then sorted into genome coordinate order and duplicate reads were flagged using Picard MarkDuplicates (http://broadinstitute.github.io/picard). Strelka^71^ 2.0.15 was used for calling single nucleotide variants and indels. Sample purity and ploidy values were estimated using ASCAT-NGS 2.1^72^. Copy number alterations after correction for estimated normal-cell contamination were inferred using ASCAT from read counts at germline heterozygous positions estimated by GATK 3.2-2 HaplotypeCaller^73^. Shallow WGS data for 75 BOs^44^ were processed with the QDNAseq package^74^ using 50-kb bins. The QDNAseq package was also used for GC-bias correction, segmentation and generation of copy number calls and used to identify homozygously deleted and amplified genes. Since the read depth was only 0.4×, mutation calls could not be generated for these samples.

### Annotation of damaged genes and OAC drivers and clonality analysis

For WGS (UoC, FHCRC) and WES (TCGA) data, SNV, indel and copy number calls were taken from the original publications or derived as described above. ANNOVAR^75^ (April 2018) and dbNSFP^76^ v3. 0 were used to annotate the effect of mutations and indels. Only SNvs and indels with damaging effects on the proteins as previously described^1^ were further retained. Briefly, these included (1) truncating (stopgain, stoploss, frameshift) mutations; (2) missense mutations predicted by at least seven methods^1^.

Copy number alteration (CNA) segments from ASCAT were intersected with the exonic coordinates of 19,641 unique human genes^1^ and a gene was considered amplified, homozygously or heterozygously deleted if at least 25% of its length overlapped with an amplified (CNA > twice sample ploidy) or homozygously (CNA = 0) or heterozygously deleted (CNA = 1) segment, respectively. Genes with at least one damaging SNV or indel as well as amplified and homozygously deleted genes were considered damaged. Genes with heterozygous deletion of one allele and at least a damaging SNV or indel in the other (double hit), were also considered damaged. Genes with only heterozygous deletions were not considered damaged. For *CDKN2A* only, *CDKN2A* silencing via methylation was also considered. Raw methylation data were processed with the minfi package in R^77^ and normalised with the BETA mixture model BMIQ of the ChAMP package^78^. *CDKN2A* was considered epigenetically silenced if the cg12840719 probe located within 1500 bp from its transcription start site^40^ had a methylation □ value ≥0.3 and its *CDKN2A* value was comparable to samples with homozygously deleted CDKN2A. The distribution of damaged genes across OAC and BO cohorts is shown in **Figure S1**. Mutated, amplified and homozygously deleted genes for the MSKCC cohort^6,37,38^ were downloaded from the cBioPortal.

Five hundred eighty out of 779 OACs with WGS or WES data (**Table S1**) had damaging alterations in *TP53* or *CDKN2A* and were further analysed to measure mutation clonality as described in^79^. Briefly, the probability of each damaging mutation to have a cancer cell fraction (CCF) from 0.01 to 1 incremented by 0.01 was calculated given the observed variant allele frequency (VAF), gene copy number status in the cancer and normal sample and sample purity. Then, the clonal probability of a *TP53* or *CDKN2A* mutation was calculated as the cumulative probability of CCF being >0.95. A damaging mutation was considered clonal if its clonal probability was >50%.

A list of 40 OAC canonical drivers was obtained from the Network of Cancer Genes (NCG7.1, http://www.network-cancer-genes.org)^1^. Additionally, 34 OAC drivers that undergo CNA were collected through manual curation of the literature. Only 54 of the resulting 74 OAC drivers were present also in the gene panel used in the MSKCC studies and these were considered for further analysis **(Table S2)**.

### Cell lines and gene expression quantification

*In vitro* experiments were carried out using the CP-A (KR-42421) Barrett’s oesophagus cells from ATCC (catalogue number CRL-4027). Cells were grown at 37°C and five per cent CO_2_ in keratinocyte serum-free medium supplemented with 50µg/ml bovine pituitary extract and 5ng/ml recombinant human EGF (Thermo Fisher). Total RNA was extracted from CP-A wild type cells and *TP53* KO clones using the Direct-zol RNA miniprep kit (ZymoResearch) and reverse transcribed using the High-capacity cDNA reverse transcription kit (Thermo Fisher). Predesigned Taqman gene expression assays for *CDKN2A* and *TP53* were used (Life Technologies, **Table S4**), while gene specific primers and probe were designed for *ACTB* (Merck, **Table S4**). Real-time quantitative PCR (rt-qPCR) was performed in duplicate using QuantiTect probe PCR mastermix (Qiagen) and repeated three times. Gene relative expression was calculated using the 2^−ΔΔCt^ method and *ACTB* as endogenous control. A pool of human RNA was used as a positive control.

### *TP53* gene editing and cell proliferation assay

To induce *TP53* KO via CRISPR-Cas9 gene editing, 3.5×10^5^ CP-A cells were co-transfected with two *TP53*-specific gRNAs (**Table S4**) and Alt-R™ S.p.Cas9-Nuclease V3 (IDT) by nucleofection using the P3 Primary Cell 4D-NucleofectorTM X Kit S (Lonza) on a 4D-Nucleofector (Lonza). After nucleofection, single cells were plated in individual wells to form clonal colonies. Genomic DNA of nucleofected colonies was extracted using PureLink Genomic DNA mini kit (Invitrogen) and regions surrounding the targeted sites were amplified from genomic DNA of nucleofected colonies using HotStartTaq Plus DNA polymerase (Qiagen) and primers including Illumina adapters (**Table S4**). Amplicons were sequenced on Illumina Novaseq using the paired-end protocol to confirm editing (BAM files: 10.5281/zenodo.12918301).

Cell proliferation of *TP53* KO and wild type CP-A cells was measured every 24 hours for three days, starting three hours after seeding the cells using CellTiter-Glo Luminescent Cell Viability Assay (Promega). Briefly, 2×10^3^ cells per well were seeded on 96-well plates in a final volume of 100μl per well. At each time point, 100μl of the CellTiter-Glo reagent was added to the wells and luminescence was measured after 30 minutes using the Infinite F200 Pro plate reader (Tecan). For all proliferation assays, two or four technical replicates per condition were measured at each time point and each measure was normalised to the average time zero measure for each condition. Each experiment was repeated three independent times. Conditions were compared using the two-tailed Student’s t-test.

### Logistic regression and survival analysis

Logistic regression with Firth bias correction^80^ was used to test the difference between two models of OAC initiation in the entire BO (P-BO and NP-BO) cohort. The first model assumed *TP53* LoF as the only driver (model 1), while the second model assumed that both *TP53* and *CDKN2A* LoF impacted on OAC initiation (model 2). The models were developed using the package logistf v1.25.0 and compared using the anova function in R. The two models were used to estimate the numbers of expected BOs that progressed to OAC according to corresponding genomic status of *TP53* and *CDKN2A*. The β coefficients for *TP53* and *CDKN2A* LoF were obtained from the regression models and the p-values were calculated using the chi-squared test. Negative or positive β coefficient values indicated cancer-protective or cancer-promoting roles, respectively. The β coefficient (β) of CDKN2A LoF in model 2 was used to estimate the odds of progression as:

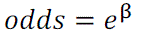

The results of the whole analysis are reported in **Table S3**.

Kaplan-Meier survival analysis was performed with survminer v.0.4.9 using the log-rank method. The analysis of the survival effect of *CDKN2A* co-damage with other 9p21 genes was performed only on 779 OAC patients with WGS or WES data as the information on the genomic alteration of all 9p21 genes was not available in the targeted re-sequencing studies. Log-rank method was used to estimate the p-values, which were then corrected for multiple hypothesis testing using the Benjamini–Hochberg method, when needed.

### RNA sequencing, gene set enrichment and immune infiltration

Paired-end RNA sequencing for OAC, P-BO and NP-BO from UoC was performed at the CRUK Cambridge Institute on Illumina platforms and quality checks were performed using FastQC. Reads were aligned using STAR with ENSEMBL gene annotation. Reads per gene were quantified using the summariseOverlaps function from the GenomicRanges package^81^. Raw read counts of 18,846 human genes shared between the UoC and TCGA cohorts were extracted from the corresponding BO and OAC RNAseq datasets. SMIXnorm v0.0.0.9^82^ was used to estimate the probability of expression of these genes across all samples. Genes with a probability of expression below 0.9 were filtered out, resulting in 16,901 retained genes in OAC, 15,134 in P-BO and 15,866 in NP-BO, respectively.

Twenty-two NP-BOs, 108 P-BOs and 337 OACs with matched genomic and transcriptomic data (**Table S1**) were divided into four groups depending on the mutation and copy number profiles of the six 9p21 genes (*KLHL9, IFNE*, *MTAP*, *CDKN2A*, *CDKN2B* and *DMRTA1*) with impact on survival. Differential gene expression analysis was performed between each of these groups and the corresponding 9p21 wild type OACs (184), P-BOs (31), and NP-BOs (6) using DESeq2 v1.38.3^58^ after correction for the batch effect with DESeqDataSetFromMatrix. Genes were ordered according to log2 fold-change values and used for pre-ranked GSEA using fgsea v1.24.0^50^ against 50 gene sets from MSigDB v7.5.1^83^ and 1,303 level 2-8 pathways from Reactome v.72^84^ containing between 10 and 500 expressed genes and excluding the ‘Disease’ hierarchical level. The resulting p-values were corrected for multiple testing in each analysis separately using the Benjamini-Hochberg method. Pathway redundancy was removed accounting for the extent of overlap between leading-edge genes, i.e. the genes that contributed the most to the enrichment. If the number of unique leading-edge genes in a pathway was higher than the shared and the unique leading-edge genes in the other pathway, the latter was removed. If the number of shared leading-edge genes between two pathways was higher than the unique leading-edge genes in both, the pathway with the higher FDR was removed. Retained processes are reported in **Table S6.**

To estimate the abundance of immune cell populations in the from bulk RNA-seq data, raw read counts of the expressed genes from 22 NP-BOs, 108 P-BOs and 337 OACs were normalized to transcripts per million (TPM) values after batch correction with ComBat-seq^85^. Resulting TPMs were used as input for ConsensusTME v0.0.1^86^ as implemented in immunedeconv v2.1.0 to estimate the normalised enrichment score (NES) using 16 oesophageal carcinoma immune signatures. To further estimate the abundance of MDSCs, two M-MDSC and G-MDSC signatures^87^ were used in ConsensusTME custom mode.

### RNAScope and Imaging Mass Cytometry

A panel of 26 antibodies targeting structural markers, immune markers, three 9p21 proteins and three RNAScope probes against *IFNE*, *IFNB1* and *PPIB* mRNAs was assembled (**Table S10**). RNAScope staining was detected using metal-tagged antibodies as previously described^88^. Sixteen of these antibodies were already metal-tagged (Standard Biotools), while eleven were carrier-free and tagged using the Maxpar X8 metal conjugation kit (Standard Biotools). The whole panel was tested in OAC FFPE sections using three dilutions ranging from 1:100 to 1:3500 and the dilution giving the highest signal-to-noise ratio was chosen for each antibody (**Table S10**).

Five μm thick sections were obtained from FFPE blocks of ten OAC patients selected based on their 9p21 gene profile (**Table S9**). Slides were incubated for one hour at 60°C, loaded on a Leica Bond autostainer (Leica Biosystems) and processed using the RNASCope LS Multiplex Fluorescent Assay following manufacturer’s instructions and *IFNE*, *IFNB1*, and *PPIB* probes at a 1:50 dilution. C2 oligos were developed with TSA-digoxinenin, C3 oligos with TSA-biotin and C1 oligos with TSA-FITC (diluted 1:200 in TSA buffer). Slides were blocked for two hours at room temperature in a Sequenza rack (Thermo Fisher Scientific). Slides were incubated overnight at 4°C with the mix of metal-conjugated antibodies, washed, and incubated with the DNA intercalator Cell-ID Intercalator-Ir (Standard Biotools). Slides were removed from the Sequenza rack, air-dried and loaded into the Hyperion Imaging System (Standard Biotools). Regions of interest (ROI) were manually selected to contain areas with tumour and immune cells by a certified pathologist (M.R.J). Regions of about 1.44 mm^2^ were laser-ablated within the pre-selected ROIs at 1 μm/pixel resolution and 400 Hz frequency.

IMC image analysis was performed using SIMPLI^89^. TIFF images for each metal-tagged antibody and DNA intercalator were obtained from the raw.txt files of the ablated regions. Pixel intensities for each channel were normalised to the 99th percentile of the intensity distribution. Background pixels of the normalised images were removed with CellProfiler4^90^ using global thresholding and processed images were verified by an expert histologist (J.S.). Single cell segmentation was performed using CellProfiler4^90^ to identify cell nucleus (DNA1 channel) and membrane (cadherin-1, pan-keratin, CD3, CD8, CD4, CD11b, CD11c, NCAM1, CD68, CD27, CD163, CD16, CD15, CD14). Obtained cells were phenotyped based on at least 10% overlap with the masks of individual cell types in the following order: (1) CD15^+^ and CD16^+^ for neutrophils; (2) NCAM1^+^ for NK cells; (3) CD11c^+^ for dendritic cells; (4) CD68^+^ for macrophages; (5) CD14^+^ for M-MDSCs; (6) CD15^+^ for G-MDSCs; (7) CD3^+^ for T cells; (8) cadherin-1 and pan-keratin for tumour cells and (9) vimentin for stromal cells. Cells with <10% overlap with any mask were left unassigned.

Unsupervised clustering was performed separately on CD3^+^ T cells and CD68^+^ macrophages using Seurat v.2.4^91^, with random seed = 123 and 0.3, 0.5, 0.7 and 0.9 cluster resolutions. Markers used for clustering were CD3, CD4, CD8, FOXP3, GzMB and Ki67 for T cells, and CD68, CD11c, HLA-DR/DP/DQ and CD163, CD11b and Ki67 for macrophages. Silhouette score of each cluster was calculated using v.2.1.6 package. The resolution with the highest median silhouette score was identified as the best clustering resolution for each cell type.

### Keratinisation causal regulatory network analysis

Causal networks linking *CDKN2A* LoF to the downregulation of keratinization were inferred using a three-step protocol modified from^92^, separately for P-BO and OAC (**Figure S4**). In the first step, co-regulated gene modules were identified using cMonkey2^93^ based on gene co-expression, proximity in the protein-protein interaction network (PPIN) and enrichment in transcription factor (TF) targets. Co-expressed genes were identified from the top 50% most variably expressed genes in P-BO and OAC after converting read counts into z-scores using DESeq2 v1.38.3^58^. Proximity in the PPIN was measured using the human weighted PPIN from STRING v11.5^94^. GO:0006355 term of Gene Ontology (release 2022-05) was used to identify 1,471 TFs. These were in turn used as input for ARACNE-AP^95^ together with P-BO and OAC gene expression data to identify TF-target pairs. cMonkey2 was run with a fixed number of iterations (n = 2000) and seed value (n = 123) for the initialization step to ensure reproducibility. The number of gene modules (k) was determined as:

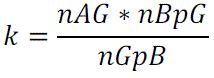

where *nAG* was the number of analysed genes, *nBpG* was the maximum number of gene modules each gene could appear in (fixed to 2), and *nGpB* was the average number of genes per gene module (fixed to 30). Identified gene modules were then filtered based on (i) co-expression quality according to the first principal component (FDR ≤0.1 and variance explained ≥0.32 for P-BO and ≥0.25 for OAC), (ii) functional enrichment in keratinisation-related genes (two-sided Fisher’s test p-value ≤0.01), (iii) enrichment in TF target genes (two-sided Fisher’s test p-value ≤0.01), (iv) correlation of TFs with gene module eigengenes, *i.e.* genes that explain the maximum expression variance. In the second step, the single.marker.analysis function of the Network Edge Orienting (NEO)^92,96^ method was used to infer causal models where *CDKN2A* LoF causally affected the expression of specific TFs, which, in turn, altered keratinisation gene modules. To assess statistical significance, the next best single marker score was defined as the log10 probability of the causal model divided by the log10 probability of the next best fitting alternative model^96^ and causal models with next best single marker score ≥0.5 were considered significant. In the third step, significant causal models were further retained if (1) TFs were differentially expressed (FDR < 0.1) in group 4 as compared to 9p21 wild type P-BOs and OACs and (2) there was significant positive correlation (R >0.5 and FDR <0.1) between TF expression and the GSEA NES score of the predicted targets in P-BOs and OACs. Finally, only TFs contributing to ≥ 30% of the significant causal models were retained. The final list of significant causal models and associated TFs is reported in **Table S11**.

## AVAILABILITY OF DATA AND MATERIALS

DNA and RNA sequence data for the University of Cambridge (UoC) cohort are available from the European Genome-phenome Archive using the following accession IDs: WGS (EGAD00001011191, EGAD00001006083), shallow WGS (EGAD00001011189), bulk RNA-sequencing (EGAD00001011190). WES for 73 TCGA OACs were downloaded from the Genomic Data Commons portal (https://portal.gdc.cancer.gov/). Mutated genes for 253 Memorial Sloan Kettering Cancer Center (MSKCC) OACs that underwent targeted re-sequencing were downloaded from the cBioPortal (https://www.cbioportal.org/). Methylation data for OACs were derived from UoC (EGAD00010001822) and TCGA (https://portal.gdc.cancer.gov/). Methylation data for BOs were derived from UoC (EGAD00010001838 and EGAD00010001972). BAM files of wild type and *TP53* edited CP-A cells are available from Zenodo (doi: 10.5281/zenodo.12918301). UoC WGS, sWGS and RNAseq data of the human patients are under controlled access by ICGC (International Cancer Consortium) due to privacy and security protection of personal data. The reasons and conditions for controlled access are described here (https://www.icgc-argo.org/page/132/data-access-and-data-use-policies-and-guidelines). The data can be accessed via the ICGC portal upon request to the ICGC Data Access Compliance Office here: https://docs.icgc-argo.org/docs/data-access/daco/applying.

## CODE AVAILABILITY

No unique code was developed for this study.

## ACKNOWLEDGEMENTS

The authors thank Patricia C. Galipeau (Fred Hutchinson Cancer Research Center, USA) for sharing somatic variant call files for the BO samples, Michele Bortolomeazzi (Deutsches Krebsforschungszentrum, Germany), Michael Pitcher and Lucia Montorsi (King’s College London, UK) for help with the IMC analysis.

## AUTHOR CONTRIBUTIONS

F.D.C. conceived and directed the study with the support of P.G.; R.C.F. recruited participants and led the OCCAMS Consortium; P.G., C.C.B., M.A., A.M., A.Z., H.M., G.D and F.D.C. analysed the data; A.A.S performed the experiments; A.B and P.B. provided some reagents and helped with cell cultures; G.D. constructed and managed the sequencing alignment and variant-calling pipelines. G.K. guided the statistical analysis. A.F. coordinated and carried out the processing of patient samples. A. F. and P.G. screened histopathological reports and identified samples; M.A., A.A.S., M.G., and E.N. performed the IMC experiments; M.R.J. and J.S. assessed tissue sections. P.G. and F.D.C. wrote the manuscript with contributions from C.C.B., M.A., A.M., A.Z., H.M., G.D. and A.F. All authors approved the manuscript.

## COMPETING INTERESTS

The authors declare no competing interests.

## OPEN ACCESS

For the purpose of Open Access, the author has applied a CC BY public copyright licence to any Author Accepted Manuscript version arising from this submission.

## FUNDING

This work was supported by Cancer Research UK [C43634/A25487 to F. D. C.] and [EDDPJT-Nov21\100010 to F. D. C], the Cancer Research UK City of London Centre [C7893/A26233 to F. D. C], Bart’s Charity and the Francis Crick Institute, which receives its core funding from Cancer Research UK (FC001002), the UK Medical Research Council (FC001002), and the Wellcome Trust (FC001002). P.B. and A.B. were supported by The Rosetrees Trust (CF2\100014).

## FULL LIST OF AUTHORS

Oesophageal Cancer Clinical and Molecular Stratification (OCCAMS) Consortium: Rebecca C. Fitzgerald^1^, Paul A.W. Edwards^1,2^, Nicola Grehan^1,5^, Barbara Nutzinger^1^, Aisling M Redmond^1^, Christine Loreno^1^, Sujath Abbas^1^, Adam Freeman^1^ Elizabeth C. Smyth^5^, Maria O’Donovan^1,3^, Ahmad Miremadi^1,3^, Shalini Malhotra^1,3^, Monika Tripathi^1,3^, Hannah Coles^1^ Curtis Millington^1^, Matthew Eldridge^2^, Maria Secrier^2^, Ginny Devonshire^1,2^, Jim Davies^4^, Charles Crichton^4^, Nick Carroll^5^, Richard H.Hardwick^5^, Peter Safranek^5^, Andrew Hindmarsh^5^, Vijayendran Sujendran^5^, Stephen J. Hayes^6,13^, Yeng Ang^6,7,26^, Andrew Sharrocks^26^, Shaun R. Preston^8^, Izhar Bagwan^8^, Vicki Save^9^, Richard J.E. Skipworth^9^, Ted R. Hupp^20^, J. Robert O’Neill^5,9,20^, Olga Tucker^10,29^, Andrew Beggs^10,25^, Philippe Taniere^10^, Sonia Puig^10^, Gianmarco Contino^10^, Timothy J. Underwood^11,12^, Robert C. Walker^11,12^, Ben L. Grace^11^, Jesper Lagergren^14,22^, James Gossage^14,21^, Andrew Davies^14,21^, Fuju Chang^14,21^, Ula Mahadeva^14^, Vicky Goh^21^, Francesca D. Ciccarelli^21^, Grant Sanders^15^, Richard Berrisford^15^, David Chan^15^, Ed Cheong^16^, Bhaskar Kumar^16^, L. Sreedharan^16^ Simon L Parsons^17^, Irshad Soomro^17^, Philip Kaye^17^, John Saunders^6, 17^, Laurence Lovat^18^, Rehan Haidry^18^, Michael Scott^19^, Sharmila Sothi^23^, Suzy Lishman^2, 24^, George B. Hanna^27^, Christopher J. Peters^27^,Krishna Moorthy^27^, Anna Grabowska^28^, Richard Turkington^30^, Damian McManus^30^, Helen Coleman^30^, Russell D Petty^31^, Freddie Bartlett^32^

^1^Early Cancer Institute, University of Cambridge, Cambridge, UK

^2^Cancer Research UK Cambridge Institute, University of Cambridge, Cambridge, UK

^3^Department of Histopathology, Addenbrooke’s Hospital, Cambridge, UK

^4^Department of Computer Science, University of Oxford, UK, OX1 3QD

^5^Cambridge University Hospitals NHS Foundation Trust, Cambridge, UK, CB2 0QQ

^6^Salford Royal NHS Foundation Trust, Salford, UK, M6 8HD

^7^Wigan and Leigh NHS Foundation Trust, Wigan, Manchester, UK, WN1 2NN

^8^Royal Surrey County Hospital NHS Foundation Trust, Guildford, UK, GU2 7XX

^9^Edinburgh Royal Infirmary, Edinburgh, UK, EH16 4SA

^10^University Hospitals Birmingham NHS Foundation Trust, Birmingham, UK, B15 2GW

^11^University Hospital Southampton NHS Foundation Trust, Southampton, UK, SO16 6YD

^12^Cancer Sciences Division, University of Southampton, Southampton, UK, SO17 1BJ

^13^Faculty of Medical and Human Sciences, University of Manchester, UK, M13 9PL

^14^Guy’s and St Thomas’s NHS Foundation Trust, London, UK, SE1 7EH

^15^Plymouth Hospitals NHS Trust, Plymouth, UK, PL6 8DH

^16^Norfolk and Norwich University Hospital NHS Foundation Trust, Norwich, UK, NR4 7UY

^17^Nottingham University Hospitals NHS Trust, Nottingham, UK, NG7 2UH

^18^University College London, London, UK, WC1E 6BT

^19^Wythenshawe Hospital, Manchester, UK, M23 9LT

^20^Edinburgh University, Edinburgh, UK, EH8 9YL

^21^Barts Cancer Institute, Queen Mary University of London, London, UK, EC1M 6BQ

^22^Karolinska Institute, Stockholm, Sweden, SE-171 77

^23^University Hospitals Coventry and Warwickshire NHS, Trust, Coventry, UK, CV2 2DX

^24^Peterborough Hospitals NHS Trust, Peterborough City Hospital, Peterborough, UK, PE3 9GZ

^25^Institute of Cancer and Genomic sciences, University of Birmingham, B15 2TT

^26^GI science centre, University of Manchester, UK, M13 9PL.

^27^Department of Surgery and Cancer, Imperial College, London, UK, W2 1NY

^28^Queen’s Medical Centre, University of Nottingham, Nottingham, UK

^29^Heart of England NHS Foundation Trust, Birmingham, UK, B9 5SS.

^30^Centre for Cancer Research and Cell Biology, Queen’s University Belfast, Northern Ireland BT7 1NN.

^31^Tayside Cancer Centre, Ninewells Hospital and Medical School, Dundee, DD1 9SY

^32^Portsmouth Hospitals NHS Trust, Portsmouth, PO6 3LY

## Notes

### Competing Interest Statement

The authors have declared no competing interest.

### Summary of Updates

In this revised version of the manuscript, we have now considered epigenetic silencing as an additional loss of function mechanism for CDKN2A and have included additional analytical and experimental validation to our hypothesis that CDKN2A early loss has a tumour suppressive role because it selects against the subsequent loss of TP53 in BO cells. We have now also reviewed the existing literature thoroughly and highlighted the controversy on the association between CKDN2A inactivation and BO progression, which justifies the need for our additional and comprehensive investigation.

